# Structure of mycobacterial ATP synthase with the TB drug bedaquiline

**DOI:** 10.1101/2020.08.06.225375

**Authors:** Hui Guo, Gautier M. Courbon, Stephanie A. Bueler, Juntao Mai, Jun Liu, John L. Rubinstein

## Abstract

Tuberculosis (TB), the leading cause of death by infectious disease worldwide, is increasingly resistant to first line antibiotics. Developed from a screen against *Mycobacterium smegmatis*, bedaquiline can sterilize even latent *M. tuberculosis* infections that may otherwise persist for decades and has become a cornerstone of treatment for multidrug resistant and extensively-drug resistant TB. Bedaquiline targets mycobacterial ATP synthase, an essential enzyme in the obligate aerobic *Mycobacterium* genus. However, how the drug binds the intact enzyme is unknown. We determined the structure of *M. smegmatis* ATP synthase with and without bedaquiline. The drug-free structure reveals hook-like extensions from the enzyme’s α subunits that inhibit ATP hydrolysis in low-energy conditions, such as during latent infections. Bedaquiline binding induces global conformational changes in ATP synthase, creating tight binding pockets at the interface of subunits a and c. These binding sites explain the drug’s structure-activity relationship and its potency as an antibiotic for TB.

## Introduction

*Mycobacterium tuberculosis* and closely related species cause the disease tuberculosis (TB), which resulted in 1.5 million deaths in 2018 alone (World Health Organization, 2019a). The same year, 0.5 million TB cases showed resistance to first line antibiotics, indicating a rise in drug resistance (World Health Organization, 2019a). The diarylquinoline drug bedaquiline (BDQ, also known as R207910, TMC207, and Sirturo) kills the mycobacteria that cause TB and other mycobacterial pathogens (Andries et al., 2005) at nanomolar concentrations. BDQ was approved by the United States Food and Drug Administration in 2012 as part of combination therapy in adults with pulmonary multi-drug resistant TB making it the first new TB drug in more than 40 years (Wellington and Hung, 2018). The World Health Organization subsequently added BDQ to its Model List of Essential Medicines (World Health Organization, 2019b). As of 2019, all seven ongoing phase III clinical trials of combination therapies for drug resistant TB include BDQ as a core component (World Health Organization, 2019a), with one trial reporting favorable outcomes in 90% of patients with extensively-drug resistant TB or complicated forms of multi-drug resistant TB (Conradie et al., 2020). It has been estimated that 80% of people with drug resistant TB would benefit from BDQ (Médecins Sans Frontières, 2019).

*M. tuberculosis* can survive low-energy conditions, allowing infections to remain dormant for extended periods and decreasing susceptibility to many antibiotics (Lu et al., 2014). BDQ can sterilize even these latent infections (Koul et al., 2008) by targeting the mycobacterial adenosine triphosphate (ATP) synthase (Andries et al., 2005), which is an essential enzyme in both *M. tuberculosis* and *M. smegmatis* (Andries et al., 2005; McNeil et al., 2020). However, affinity measurements between the drug and the isolated c subunit of ATP synthase, which was identified as part of its binding site, showed only micromolar affinity (Haagsma et al., 2011; Koul et al., 2007) suggesting that binding of the drug to the intact enzyme involves additional subunits. ATP synthases use energy stored in a transmembrane proton motive force established during respiration to synthesize ATP from adenosine diphosphate (ADP) and inorganic phosphate. In the absence of a proton motive force during low-energy conditions, the ATP synthase must be prevented from running in reverse and hydrolyzing the cell’s supply of ATP. How this inhibition of ATP hydrolysis occurs in mycobacteria is unclear. In order to understand the structural basis by which ATP hydrolysis by ATP synthase is inhibited in mycobacteria, and how BDQ targets mycobacterial ATP synthase, we isolated the *M. smegmatis* ATP synthase and used electron cryomicroscopy (cryoEM) to determine its structure with and without BDQ bound.

## Results

### Overall structure of mycobacterial ATP synthase includes an unusual peripheral stalk

Similar to other bacterial ATP synthases (Guo et al., 2019; Sobti et al., 2020), the mycobacterial ATP synthase consists of a soluble cytosolic F_1_ region and a membrane-embedded F_O_ region (Lu et al., 2014). To isolate endogenous ATP synthase from *M. smegmatis*, we introduced a sequence into the chromosomal DNA encoding 3×FLAG affinity tags at the C termini of the β subunits in the F_1_ region. The resulting strain was cultured, its membranes harvested, and the ATP synthase purified by affinity chromatography. CryoEM of the preparation (Fig. S1 and S2) yielded three three-dimensional (3D) maps, corresponding to the main rotational states of the enzyme (Guo et al., 2019; Zhao et al., 2015), at nominal resolutions of 3.4 to 3.7 Å. Focused refinement of the F_O_ region (Fig. S1 and S2) produced a map at 3.5 Å resolution (Fig. 1A, *left*). Together, these maps allowed construction of an atomic model for almost all of the complex (Fig. 1A, *right*, Fig. S3, Tables S1-S5).

**Figure 1.**
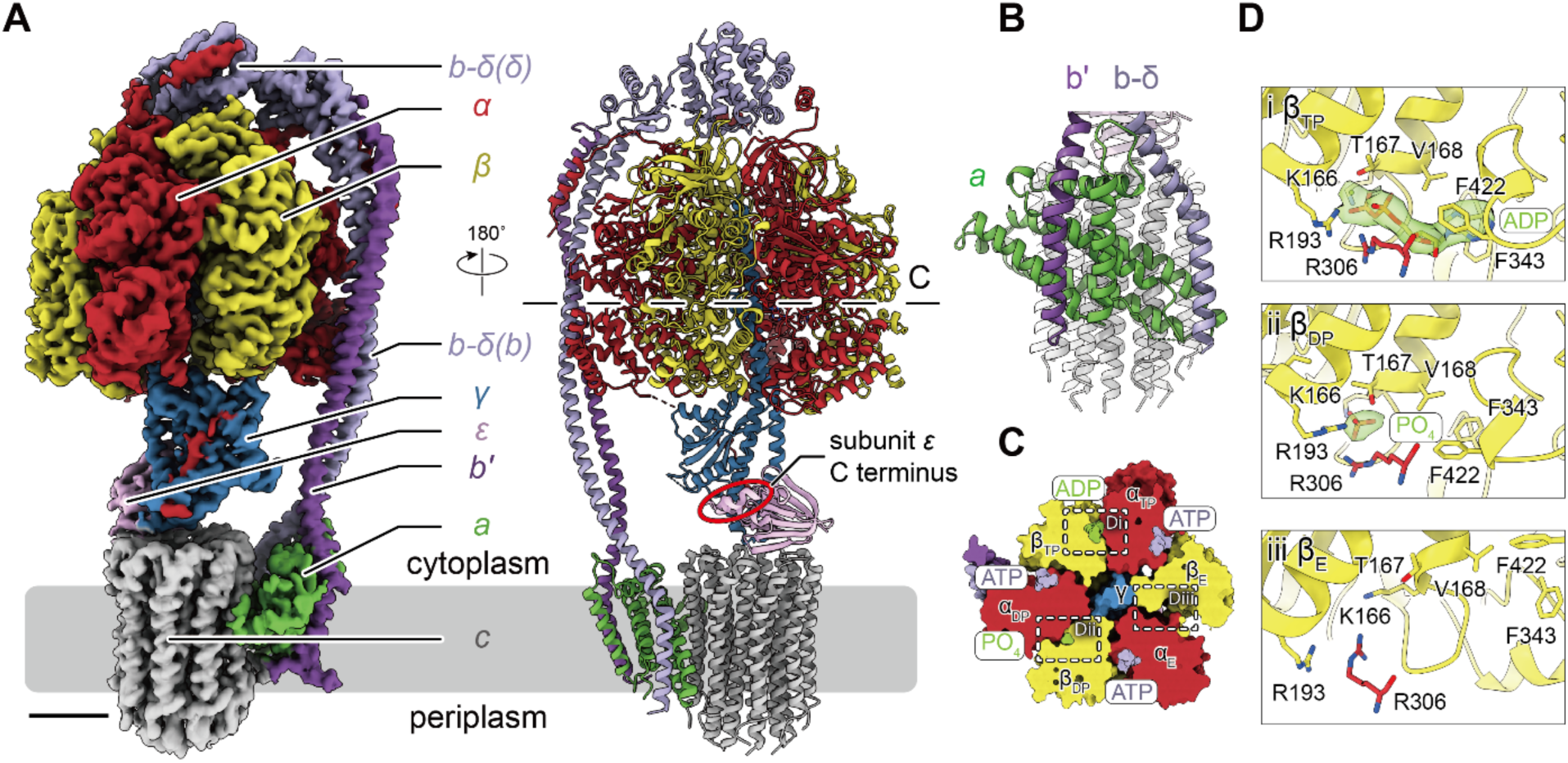
Structure of the mycobacterial ATP synthase. **A**, Composite cryoEM density map (*left*) and atomic model (*right*). Scale bar, 25 Å. The *red oval* indicates that subunit ε is in a non-inhibitory ‘down’ conformation. **B**, Interaction of subunit b*′* and b-δ with subunit a in the membrane. **C**, Cross section through the F_1_ region of the model showing nucleotide content consistent with an uninhibited enzyme. **D**, Close-up view of nucleotide content in the catalytic sites.

The *M. smegmatis* F_O_ region contains nine copies of subunit c (Fig. 1A and B, *grey*), similar to the isolated c ring from *M. phlei* (Preiss et al., 2015). The proton conducting subunit a (Fig. 1A and B, *green*) has the same fold found in the mammalian (Zhou et al., 2015), yeast (Guo et al., 2017), algal (Klusch et al., 2017), and chloroplast (Hahn et al., 2018) ATP synthases and aligns well with the *Bacillus* PS3 and *E. coli* subunits (Fig. S4) (Guo et al., 2019; Sobti et al., 2020). During ATP synthesis, the transmembrane proton motive force drives protons through the F_O_ region, inducing rotation of the γεc_9_ rotor subcomplex (Junge et al., 1997; Vik and Antonio, 1994). This rotation of γ within the catalytic α_3_β_3_ subcomplex of the F_1_ region leads to synthesis of ATP from ADP and inorganic phosphate. The α_3_β_3_ subcomplex is prevented from rotating in sympathy with the rotor by a peripheral stalk, which consists of two copies of subunit b and subunit δ in most bacteria (Guo et al., 2019; Sobti et al., 2016) but is formed from a b-δ fusion protein and subunit b*′* in mycobacteria (Lu et al., 2014). The structure shows that the transmembrane α helices of subunit b*′* and subunit b-δ sandwich the N-terminal α helix of subunit a, replacing the two identical b subunits found in the canonical bacterial ATP synthase (Fig. 1B) (Sobti et al., 2016). Subunit b*′* is packed tightly against the second and third transmembrane α helices of subunit a while subunit b-δ forms fewer contacts. These interactions explain the observation that subunit b-δ is responsible for mediating attachment with the F_1_ region while subunit b*′* is more important for binding the F_O_ region (Gajadeera and Weber, 2013). The δ region of the b-δ fusion subunit differs dramatically from the canonical bacterial ATP synthase δ subunit (Fig. S5) (Guo et al., 2019; Sobti et al., 2016).

### Hook-like structures from the α subunits prevent ATP hydrolysis

The F_1_ regions of ATP synthases have three catalytic nucleotide-binding sites, primarily within the β subunits, and three non-catalytic sites, primarily within the α subunits (Fig. 1C and D) (Abrahams et al., 1994). These catalytic sites are known as β_TP_ (indicating ‘ATP-bound’), β_DP_ (indicating ‘ADP-bound’), and β_E_ (indicating ‘Empty’), based on their expected nucleotide content. Because ATP synthases can function as ATPases that deplete cellular energy reservoirs, these enzymes require a mechanism to inhibit ATP hydrolysis in the absence of a transmembrane proton motive force. Inhibition of ATP hydrolysis is thought to be particularly important in pathogenic mycobacteria due to low-energy challenges faced within hosts (Lu et al., 2014). In other bacteria (Cingolani and Duncan, 2011; Guo et al., 2019; Shirakihara et al., 2015), subunit ε blocks ATP hydrolysis by adopting an inhibitory ‘up’ conformation where its C-terminal α helix inserts into a catalytic α/β interface. However, as seen previously (Zhang et al., 2019), the purified mycobacterial ATP synthase has only negligible ATP hydrolysis activity (Fig. 2A, *left*) even though subunit ε adopts a non-inhibitory ‘down’ conformation (Fig. 1A, *right, red oval*). The structure of the *M. smegmatis* F_1_ region purified in the presence of ADP showed MgADP in both the β_TP_ and β_DP_ sites and a weak density, consistent with phosphate or sulfate, in the β_E_ site (Zhang et al., 2019). This unusual nucleotide occupancy was suggested as an explanation for inhibition of ATP hydrolysis activity (Zhang et al., 2019). For the present study no exogenous nucleotide was added. In this structure (Fig. 1C and D), only the β_TP_ site has density for ADP, while the β_E_ site is empty and density consistent with phosphate or sulfate is found in the β_DP_ site, indicating that a different explanation for inhibition of hydrolysis must exist.

**Figure 2.**
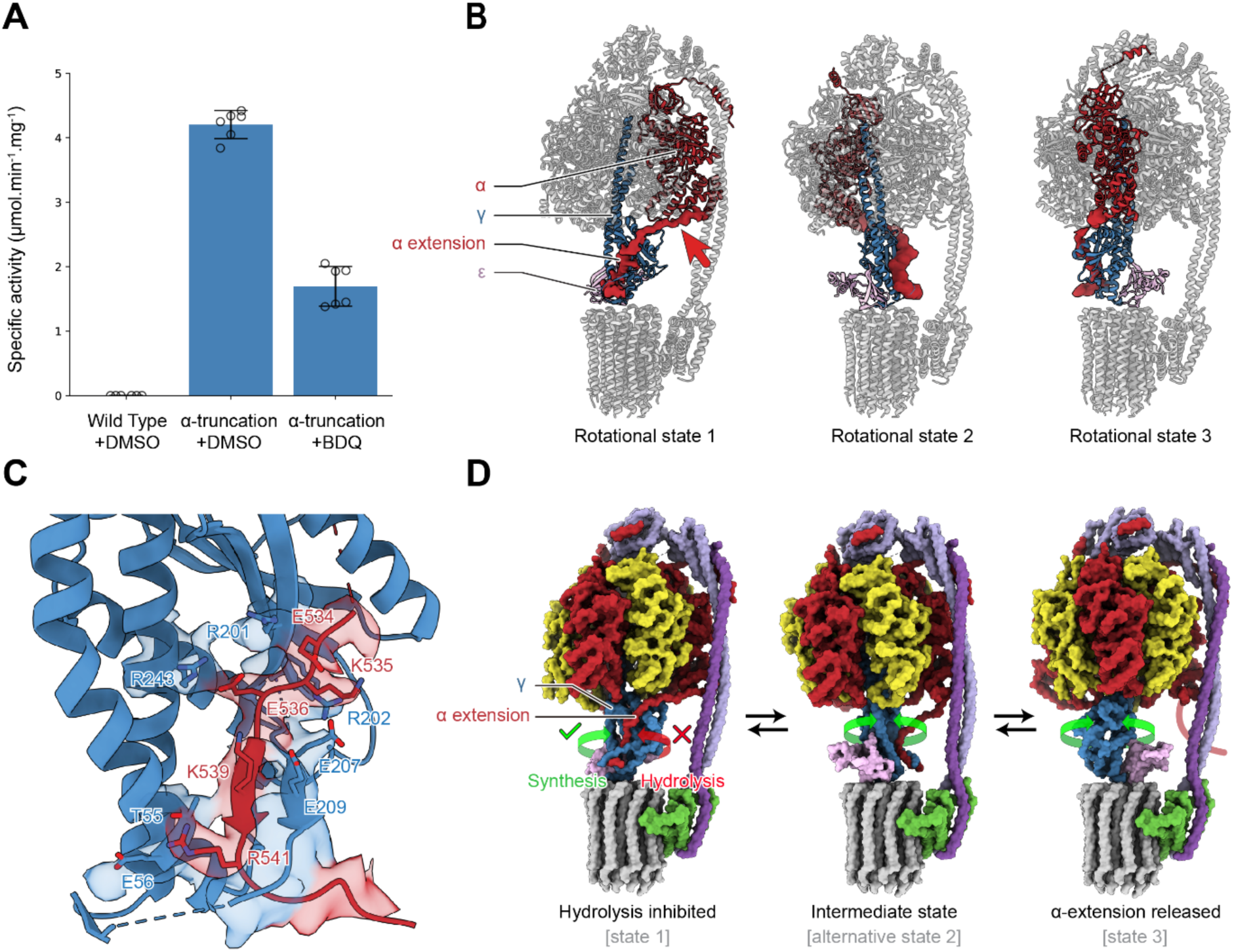
Autoinhibition of ATP hydrolysis. **A**, ATP hydrolysis is inhibited in intact mycobacterial ATP synthase, but inhibition is released by truncation of the α extension at Ser518. ATP hydrolysis in the enzyme with truncated α extension is inhibited ∼60% by addition of BDQ. Error bars, ±SD (N=6, two biological replicates with three technical replicates each). **B**, Each rotational state shows one α extension bound to the γ subunit. **C**, The α extension interacts with subunit γ by forming a two-stranded parallel β sheet. Ser518 of subunit α is indicated with a *red arrow*. **D**, Rotation in the ATP hydrolysis direction is inhibited by binding of the α extension but rotation in the ATP synthesis direction releases the α extension (see also Video 1).

Instead, the mycobacterial ATP synthase appears to have a unique mechanism for inhibiting ATP hydrolysis (Video 1). Mycobacterial α subunits possess ∼35 residue C-terminal extensions not found in other ATP synthases (Zhang et al., 2019). These extensions were previously proposed to form α helices involved in inhibition (Ragunathan et al., 2017), but this model was questioned because the extensions were not observed in a crystal structure of the F_1_ region and were predicted to be disordered (Zhang et al., 2019). However, in the present study a single α extension is resolved in the map for each rotational state, forming intimate contacts with the γ subunit (Fig. 2B, *red*). The remaining two α extensions are not resolved, suggesting that they are disordered when not bound to subunit γ. An evolutionarily conserved region of the α extension (Zhang et al., 2019) forms a β strand that creates a parallel β sheet with a β strand from subunit γ and possesses multiple charged residues that appear to stabilize this α/γ interaction (Fig. 2C). Consequently, the C terminus of the α extension is trapped under the β strand from subunit γ, forming the hook of a rachet that only allows unidirectional rotation of subunit γ in the ATP synthesis direction and preventing ATP hydrolysis.

To test this model of inhibition, an *M. smegmatis* strain was constructed with a 3×FLAG tag inserted after residue Ser518 of subunit α (Fig. 2B, *red arrow*). This construct truncates the α subunits, removing the C-terminal extensions. The resulting complex had strong ATP hydrolysis activity (Fig. 2A, *middle*), which could be inhibited by ∼60% upon addition of bedaquiline (BDQ; also known as R207910, TMC207, and Sirturo) (Fig. 2A, *right*). Rotational state 2 also revealed an alternative conformation with the α extension from a different α subunit bent in the opposite direction to attach to the central rotor (Fig. 2D, *centre*, Fig. S6). The existence of this conformation suggests the sequence of events that occurs as the α extension binds subunit γ to block ATP hydrolysis or is released from subunit γ during ATP synthesis (Fig. 2D, Video 1). While binding of an α extension prevents rotation in the ATP hydrolysis direction (Fig. 2D, *left*), rotation in the ATP synthesis direction causes the α extension to fold back on itself, as seen in the alternative conformation of rotational State 2 (Fig. 2D, *middle*). Further rotation in the ATP synthesis direction pulls the C terminus of the α extension out from under subunit γ, breaking the interaction between the two proteins (Fig. 2D, *right*).

### Bedaquiline binds three distinct sites in ATP synthase

*In vitro* selection and sequencing of three resistance mutants established that the mechanism of action of BDQ (Fig. 3A) involves the mycobacterial ATP synthase c subunits (Andries et al., 2005; Koul et al., 2007). However, only micromolar affinity was measured between BDQ and recombinant subunit c (Haagsma et al., 2011; Koul et al., 2007), which typically assembles into detergent-resistant rings upon expression (Preiss et al., 2015). This micromolar binding affinity is inconsistent with the drug’s nanomolar inhibition of mycobacterial ATP synthesis (Koul et al., 2007; Preiss et al., 2015) and nanomolar inhibition of *M. tuberculosis* growth (Andries et al., 2005). Consequently, it has been suggested that subunit a in the F_O_ region also contributes to BDQ binding in the intact complex *in vivo* (Haagsma et al., 2011). To determine how BDQ interacts with intact mycobacterial ATP synthase, the enzyme was incubated with 200 µM drug and again subjected to cryoEM structure determination. This BDQ concentration corresponds to a ∼40-fold molar excess over the enzyme but only a ∼4-fold molar excess relative to the c subunits in the complex. Image analysis produced maps of the three rotational states at nominal resolutions of 3.2 to 3.4 Å with focused refinement of the F_O_ region providing a map at 3.4 Å resolution (Fig. S1 and S2). The maps revealed seven BDQ molecules, one interacting with each c subunit in the ring except for the two c subunits occluded by subunit a (Fig. 3B, Video 2). No density was found to support the proposed binding of BDQ to subunit ε (Kundu et al., 2016). Five of the binding sites, which we refer to as ‘c-only sites’, correspond to sites described previously from a structure of isolated *M. phlei* c ring crystallized in the presence of millimolar BDQ (Preiss et al., 2015) (Fig. 3B-C, *yellow*). Two previously unobserved sites also involve interactions with subunit a. Considering the direction of rotation of the c ring during ATP synthesis (Fig. 3B, *purple arrow*), one of these sites, which we name the ‘leading site’(Fig. 3B-C, *pink*) involves a c subunit that has just interacted with subunit a and picked up a proton from the periplasmic half channel. The other site, which we designate the ‘lagging site’ (Fig. 3B-C, *blue*) is poised to interact with subunit a to deposit a proton into the cytoplasmic half channel.

**Figure 3.**
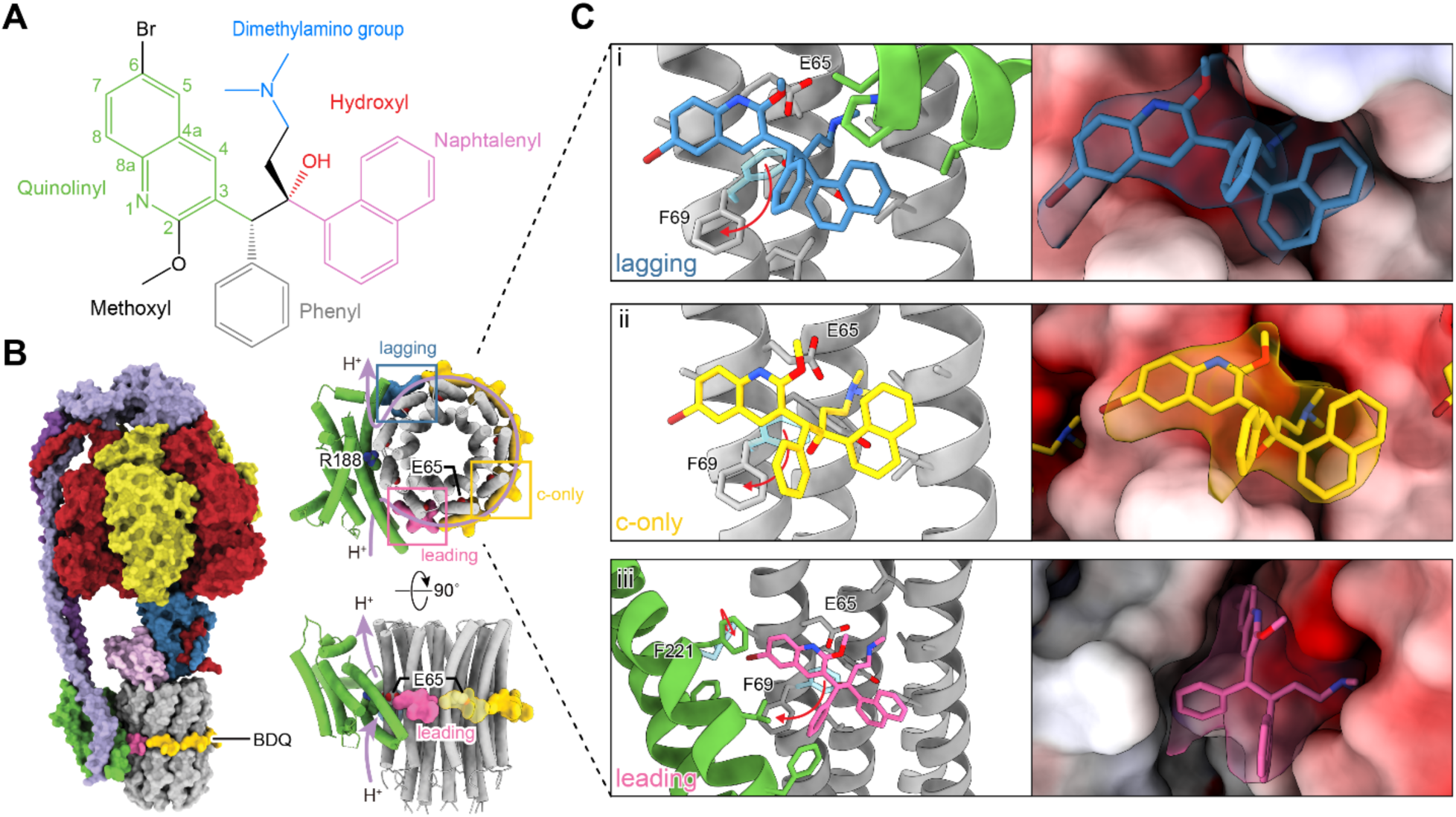
Inhibition of mycobacterial ATP synthase by BDQ. **A**, Structure of BDQ. **B**, In saturating conditions, BDQ binds seven sites in the mycobacterial ATP synthase: five ‘c-only’ sites (*yellow*), a leading site (*pink*), and a lagging site (*blue*). The direction of proton flow and c ring rotation are indicated (*purple arrow*). **C**, Atomic model (*left*) and surface representation (*right*) of the lagging (*top*), c-only (*middle*), and leading (*bottom*) BDQ binding sites. The *red arrows* indicate movement of residues upon BDQ binding (F69 in subunit c and F221 in subunit a), with their position in the drug-free structures shown in transparent light blue.

In the five c-only sites, the dimethylamino group of BDQ penetrates into the ion-binding pocket of the subunit, interacting with the carboxyl group of the proton-carrying Glu65 residue, as described previously (Preiss et al., 2015) (Fig. 3Cii). This interaction is consistent with the finding that the tertiary amine of BDQ is necessary for the drug’s antimycobacterial activity (Lounis et al., 2010). Also seen previously (Preiss et al., 2015), Phe69 of subunit c moves from its orientation in the drug-free structure away from BDQ in the drug-bound structure (Fig. 3Cii, *red arrow*). This movement avoids a clash with BDQ’s hydroxyl group and provides what has been called a ‘hydrophobic platform’ for the quinoline moiety (Preiss et al., 2015). The remainder of the c-only binding pocket is formed primarily from Ala66 and Tyr68 from the same c subunit as Glu65 as well as Leu63, Ala66, Ala67, Ile70 from the adjacent c subunit. BDQ’s bromine group remains accessible on the surface of the complex, supporting the reliability of affinity measurements between BDQ and the c ring where the drug was tethered with a linker at position 6 of the quinolinyl group (Fig. 3A, *green*) (Haagsma et al., 2011; Koul et al., 2007). An explanation for BDQ’s activity at low concentration can be found in the binding sites that only exist in the intact ATP synthase. The lagging drug binding site is clockwise from subunit a when viewed from the F_1_ region toward the F_O_ region (Fig. 3B, *blue*). This binding pocket is formed from the c subunit residues described above, as well as Leu70, Pro172, Ile173, and Val176 from subunit a. In the lagging site, Phe69 of subunit c undergoes the same conformational change seen in the c-only sites. The additional interactions provided by subunit a in the lagging site produce a deeper binding pocket than the c-only pocket, which increases protein-drug Van der Waals contacts (Fig. 3Ci) but does not appear to induce additional side chain rearrangement in the protein. The leading site, which is likely the most important for BDQ’s *in vivo* activity, is counter-clockwise from subunit a when the complex is viewed from F_1_ toward F_O_ (Fig. 3B, *pink*). In addition to the c subunit residues found in the c-only and lagging sites, the leading site binding pocket comprises residues Phe213, Pro214, Val217, Trp218, and Phe221 from subunit a. These residues create a particularly deep binding pocket that fits BDQ with extensive contacts (Fig. 3Ciii). The bromine group of BDQ interacts with Phe221of subunit a in the leading site, with the side chain undergoing a conformational change to accommodate drug binding (Fig. 3Ciii, *red arrow*). This interaction explains structure-activity relationship studies of the diarylquinoline series that led to BDQ, where the bromine increased antimycobacterial activity (Guillemont et al., 2011). It is also consistent with the finding that halogens in ligands have a high propensity to interact with hydrophobic amino acids (Kortagere et al., 2008), with interactions between bromine and Phe residues common in drug binding (Sirimulla et al., 2013).

### The leading and lagging binding sites explain bedaquiline’s potency

Binding of BDQ to the leading and lagging sites induces a global conformational change in the enzyme, trapping the rotor in an intermediate position between the rotary states described above (Fig. 4A, Video 2). The direction of rotation in each state can be determined by inspecting the F_1_ region, and amounts to a 28° rotation in the synthesis direction for State 1, 25° in the synthesis direction for State 2, and 8° in the hydrolysis direction for State 3 (Fig. 4B). This rotation is needed to produce both the leading and lagging binding sites. The extensive contact between the drug and protein suggests that the leading and lagging sites may have a higher affinity than the c-only sites. To test this hypothesis, which would explain the activity of BDQ at low concentration, we attempted to remove weakly-bound BDQ by washing the drug-bound complex in a desalting column with a drug-free buffer. CryoEM of the resulting preparation gave maps of the three rotational states at nominal resolutions of 3.3 to 3.5 Å resolution and the F_O_ region at 3.7 Å resolution (Fig. S1 and S2). The conformations of the structures matched the drug-bound state rather than the drug-free state (Fig. S7). However, in the washed structure, density for BDQ in the c-only sites is extremely weak (Fig. 4C and D, *yellow*) indicating that most BDQ has been removed from these sites. Further, density for the Phe69 residues in the c-only sites more closely resembles the density in the drug-free state than the BDQ-saturated state (Fig. 4D, *red arrow*). In contrast, strong density for BDQ is found in the leading site (Fig. 4C and D, *pink*), with weaker density for BDQ in the lagging site (Fig. 4C and D, *blue*). Density for the Phe69 residues in the leading and lagging sites resembles the drug-saturated state. The clarity of Tyr68 and Phe64 in the maps support the ability to resolve side-chain detail in all three sites (Fig. 4D). Together, these observations suggest that the leading site binds BDQ the most tightly while the c-only sites bind it with lower affinity. The binding of BDQ in the leading and lagging sites, which are not present in the isolated c ring, explains the potent antimycobacterial activity of BDQ at low concentration.

**Figure 4.**
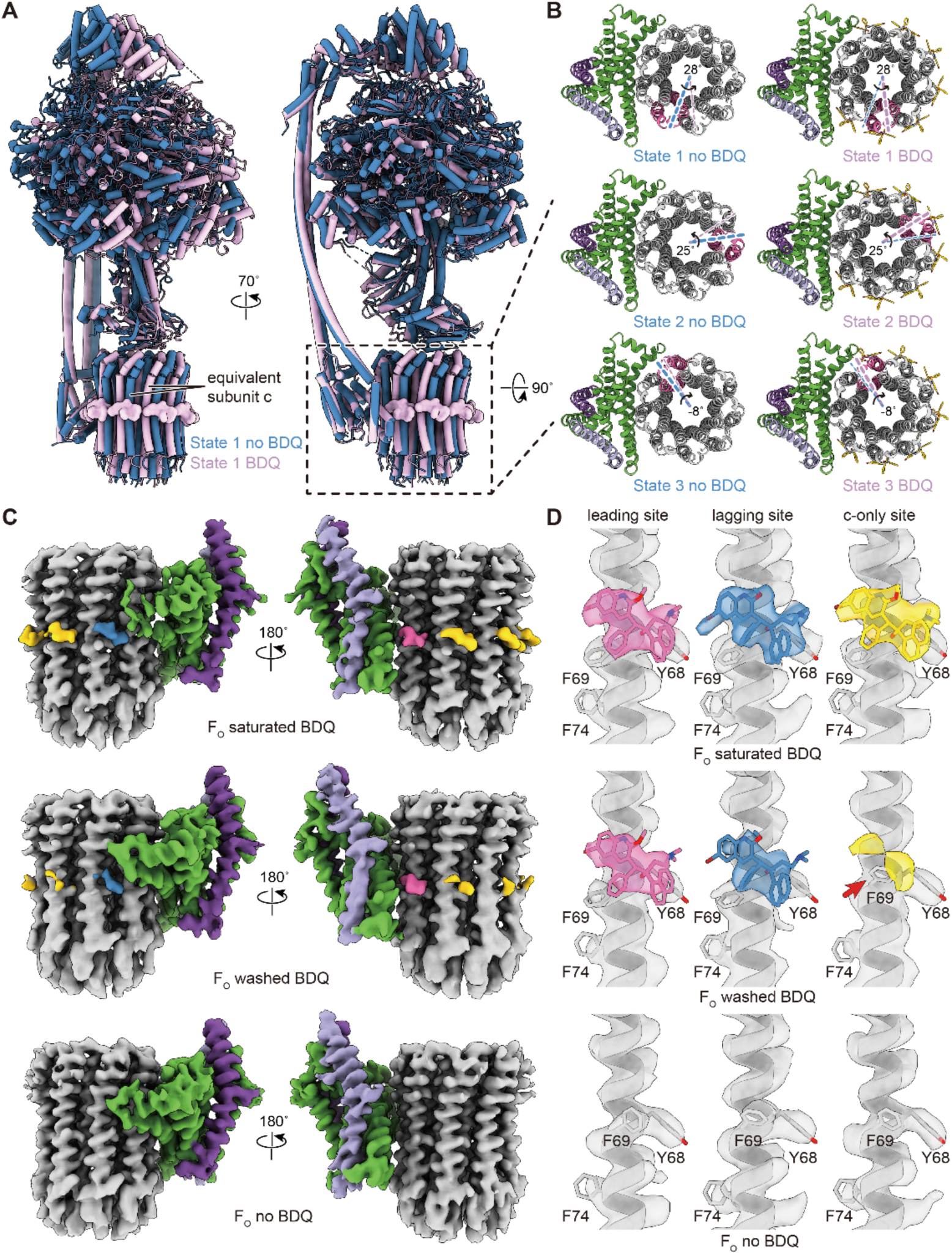
Large scale conformational changes and high-affinity binding of BDQ. **A**, Shown for rotational state 1, addition of BDQ induces rotation of the rotor to form drug binding sites. **B**, The induced conformational changes from BDQ binding corresponds to a 28° rotation in the ATP synthase direction for rotational state 1, 25° in the ATP synthase direction for rotational state 2, and 8° in the ATP hydrolysis direction for rotational state 3. **C**, Comparison to the BDQ-saturated (*top*) and BDQ-free (*bottom*) structures shows that removal of weakly-bound BDQ by washing (*middle*) leads to loss of density at the c-only sites, indicating a lower affinity for these sites. **D**, Map densities for F69 in the c-only sites of the washed complex resemble the BDQ-free state (*red arrow*) while in the leading- and lagging-sites they match the drug-bound state. Density for Y68 and F74 support the map resolution in all of the sites. (see also Video 2).

## Discussion

The newly-identified BDQ binding sites are formed by induced fit, with ATP synthase undergoing rotation of its c ring away from the position found in the resting drug-free structure. Rotation of the ring allows Glu65 of subunit c to interact with the drug’s dimethylamino group while simultaneously creating close but distinct contacts between BDQ and subunit a in the leading and lagging sites. The sequences of mycobacterial ATP synthase subunits a and c differ from the human ATP synthase in these binding pockets, explaining BDQ’s specificity for mycobacteria (Haagsma et al., 2009; Preiss et al., 2015). Consequently, in addition to revealing the unique mechanism by which the ATP synthase prevents ATP hydrolysis in the low-energy conditions faced by mycobacterial pathogens, the structures provide information that can be used for further optimization of BDQ. They also show that as the c ring rotates it creates potential drug binding sites of varying size that bridge subunits c and a, which could be targeted by other compounds to inhibit ATP synthesis and kill mycobacteria.

## Acknowledgements

We thank Voula Kanelis for suggestions on the manuscript. HG was supported by an Ontario Graduate Scholarship for International Students. JLR was supported by the Canada Research Chairs program. This work was supported by Canadian Institutes of Health Research grant PJT162186 (JLR) and PJT156261 (JL). CryoEM data was collected at the Toronto High-Resolution High-Throughput cryoEM facility, supported by the Canada Foundation for Innovation and Ontario Research Fund.

## Author contributions

JLR conceived the project and supervised the research. JL advised on *M. smegmatis* cell culture and chromosomal tagging strategies. SAB prepared the a 3×FLAG ORBIT payload plasmid and an *M. smegmatis* strain used in the initial stages of the project with help from JM. GMC prepared the *M. smegmatis* strains used for structure determination and enzyme assays. GMC and HG purified the protein, HG performed the cryoEM and image analysis, and GMC and HG constructed the atomic models. JLR, HG, and GMC wrote the manuscript and prepared figures with input from the other authors.

## Data and materials availability

CryoEM maps are deposited in the Electron Microscopy Data Bank under accession numbers EMD-22311 to 22322. Atomic models are deposited in the Protein Data Bank under accession numbers7JG5, 7JG6, 7JG7, 7JG8, 7JG9, 7JGA, 7JGB, 7JGC. Strains and plasmids are available from the corresponding author.

## Methods

### Plasmid and strains

*M. smegmatis* strains were generated by the ORBIT (oligonucleotide-mediated recombineering followed by Bxb integrase targeting) method (Murphy et al., 2018), which requires a plasmid with the Che9c phage RecT annealase and Bxb1 integrase, a payload plasmid with the desired insert, and an oligonucleotide that guides integration of the payload into the chromosomal DNA. The FLAG-His sequence in payload plasmid pKM491 (Addgene) was replaced with sequence for a 3×FLAG tag by Gibson Assembly (New England Biolabs) to create payload plasmid pSAB41. *M. smegmatis* strains were then constructed by transforming strain MC^2^155 bearing plasmid pKM444, which encodes the Che9c annealase and Bxb1 integrase, with plasmid pSAB41 and the targeting oligonucleotide. Strain GMC_MSM1 was generated with a 3×FLAG tag at the C terminus of the β subunits with oligonucleotide 5*′*-CGACGGCGACGATCTCGACGTTCAGATCAGCCACACCACACCACCTTTCGGACGCTCAAGGACATGGCAATCAGGTTTGTACCGTACACCACTGAGACCGCGGTGGTTGACCAGACAAACCCAGCTTGGCGCCGAGGCTCTCGGCCTTCTTCGCCAGGTCGTCCAGACCACCGATCAGGAAGAACGCCTGC-3*′*. Strain GMC_MSM2, with the α subunits truncated after Ser 518 followed by a C-terminal 3×FLAG tag, was generated with oligonucleotide 5*′*-TGCGGACCTTGACGGATTCCTTCTCCAGGTCCTCGGGATCGAGGGCCTCGGCGTTCTCGGAGACGACCACGGTTTGTACCGTACACCACTGAGACCGCGGTGGTTGACCAGACAAACCCGAGCTGCCGTCAGAGGCCTGGAAGCCCTTCTTGAATTCGTTGATGACCGAGACCAGCTTCTCCTCGGCT-3*′*. Transformants were selected with hygromycin (50 μg/ml) and correct insertion of the 3×FLAG sequence was confirmed by colony PCR.

### Bacterial growth and protein purification

*M. smegmatis* was grown in Middlebrook 7H9 broth supplemented with 0.8 g/L NaCl, 2 g/L dextrose, 10 g/L tryptone, and 0.05% (v/v) Tween 80. Cells were harvested by centrifugation and resuspended in 4 mL/g lysis buffer (50 mM Tris-HCl pH 7.5, 150 mM NaCl, 5 mM MgSO_4_, 5 mM 6-aminocaproic acid, 5 mM benzamidine, 1 mM PMSF). The suspension was filtered through Miracloth (Millipore), before 3 to 4 passages through an EmulsiFlex–C3 High Pressure Homogenizer (Avestin) at 20 kpsi. Unbroken cells and debris were removed by centrifugation at 39,000 g for 30 min. Membranes were collected by centrifugation at 257,000 g for 90 min and resuspended in buffer (50 mM Tris-HCl pH 7.4, 15% [v/v] glycerol, 5 mM MgSO_4_, 150 mM NaCl, 5 mM 6-aminocaproic acid, 5 mM benzamidine, 1 mM PMSF) prior to storage at -80 °C. For protein purification, membranes were thawed and solubilized with 1% (w/v) dodecyl-β-D-maltoside (DDM) (Anatrace) and insoluble material was removed by centrifugation at 257,000 g for 60 min. Solubilized protein was then loaded onto 2 mL M2 affinity matrix (Sigma) previously equilibrated with wash buffer (50 mM Tris-HCl pH 7.4, 15% [v/v] glycerol, 150 mM NaCl, 5 mM 6-aminocaproic acid, 5 mM benzamidine, 0.05% DDM [w/v]) and the ATP synthase eluted with 150 µg/ml 3×FLAG peptide in wash buffer. The sample was concentrated and loaded onto a Superose 6 Increase 10/300 column (GE Healthcare) equilibrated with gel filtration buffer (50 mM Tris-HCl pH 7.4, 15% [v/v] glycerol, 150 mM NaCl, 0.05% DDM [w/v]). Eluted ATP synthase was pooled and concentrated to ∼6 mg/ml and stored at −80 °C. ATP hydrolysis activity was measured as described previously (Kornberg and Pricer, 1951; Vasanthakumar et al., 2019) in ATPase buffer (50 mM Tris-HCl pH 7.4, 150 mM NaCl, 10% [v/v] glycerol, 5 mM MgCl_2_, 0.2 mM NADH, 2 mM ATP, 1 mM phosphoenol pyruvate, 3.2 units pyruvate kinase, 8 units lactate dehydrogenase, 0.05% [w/v] DDM). Inhibition with BDQ (AmBeed) was assayed at 2 µM.

### CryoEM

Holey gold grids (Meyerson et al., 2014; Russo and Passmore, 2014) with a regular arrays of ∼2 µm holes were nanofabricated as described previously (Marr et al., 2014). CryoEM specimens of ATP synthase without BDQ were prepared following removal of glycerol with a Zeba Spin desalting column (Thermo Fisher Scientific) with protein (2.5 μL) applied to grids that had been glow-discharged in air for 2 min. Grids were then blotted in a Vitrobot Mark III (Thermo Fisher Scientific) for 26 s at 4 °C and ∼100% humidity before freezing in a liquid ethane/propane mixture (Tivol et al., 2008). Specimens of BDQ-saturated ATP synthase were prepared by mixing 1 μL of 2.5 mM BDQ in DMSO with 12 μL of 5 mg/ml ATP synthase and incubating on ice for 1 h. Immediately before freezing, glycerol was removed with a Zeba Spin desalting column equilibrated with buffer containing 0.2 mM BDQ (final DMSO concentration 2%). Specimens of ATP synthase with weakly-bound BDQ washed away were prepared by excluding BDQ from the final desalting step with the Zeba Spin column.

### Data collection

Cryo-EM images were collected at 300 kV with a Titan Krios G3 electron microscope equipped with a Falcon 4 camera (Thermo Fisher Scientific). Automated data collection was done with the *EPU* software package. For the drug-free dataset, 7,691 movies consisting of 30 exposure fractions were collected at a nominal magnification of 75,000×, corresponding to a calibrated pixel size of 1.03 Å. The camera exposure rate and the total exposure for the specimen were 5.0 e^-^/pix/s and ∼45 e^-^/Å^2^, respectively. For the BDQ-saturated dataset, 7,589 movies consisting of 29 exposure fractions were collected at the same magnification. The camera exposure rate and the total exposure of the specimen were 4.9 e^-^/pix/s and ∼41 e^-^/ Å^2^, respectively. The BDQ-washed dataset consisted of 4,962 movies at the same conditions as the BDQ-saturated dataset.

### Image analysis

Except where noted, image analysis was performed with *cryoSPARC* v2 (Punjani et al., 2017). Movie frames were aligned with *MotionCor2* (Zheng et al., 2017) using a 7×7 grid and contrast transfer function (CTF) parameters were estimated in patches. Templates for particle selection were generated by 2D classification of manually-selected particle images and datasets of particle images were extracted in 320×320 pixel boxes. This process provided 1,435,679 particle images for the drug-free condition, 1,825,672 particle images for the BDQ-saturated condition, and 1,865,961 particle images for the BDQ-washed condition. Datasets were cleaned with two rounds of 2D classification, reducing their size to 708,053, 797,936, and 608,800 particle images for the three conditions, respectively. After ab initio 3D classification and heterogeneous refinement, three classes corresponding to the main rotational states of the complex were identified in each dataset. Classes were refined with non-uniform refinement (Punjani et al., 2019), individual particle defocus parameters were estimated, and maps were refined again with non-uniform refinement. Image parameters were then converted to *Relion 3*.*0* (Scheres, 2012) .star file format with the *pyem* package (DOI: <u>10.5281/zenodo.3576630</u>) and beam-induced motion was corrected for individual particles in *Relion* (Zivanov et al., 2019). These images were imported back to *cryoSPARC* v2, CTF parameters were re-estimated, and non-uniform refinement and several rounds of ab initio 3D classification followed by heterogeneous refinement were performed to remove poorly behaved particle images from each class. A final round of non-uniform refinement produced 3D maps of the intact complex in the different rotational states. For the drug-free condition the three classes contained 152,372, 46,960, and 120,424 particle images and reached 3.4 Å, 3.7 Å and 3.5 Å resolution, respectively. Masked local refinement of the F_O_ region with 319,756 particle images from all three classes yielded a map at 3.5 Å resolution. For the BDQ-saturated condition the three classes contained 112,421, 70,566, and 155,488 particle images and reached 3.3 Å, 3.4 Å and 3.2 Å resolution, respectively. Local refinement of the F_O_ region of State 3 yielded a map at 3.4 Å resolution. For the BDQ-washed condition the three rotational state classes contained 97,912, 69,407, and 127,165 particle images and reached 3.3 Å, 3.5 Å and 3.3 Å resolution, respectively. Masked local refinement of the F_O_ region with 294,484 particle images from all three classes yielded a map at 3.7 Å resolution. All the maps were locally sharpened with LocalDeblur (Ramírez-Aportela et al., 2020). For illustration purposes, composite maps of some of the rotational states were generated by combining the F_1_ region of the intact maps with the F_O_ region of the maps from masked refinement using *UCSF Chimera* (Pettersen et al., 2004) and *relion_image_handler*.

### Atomic model building and refinement

The State 1 map for the BDQ-free specimen and the State 3 map for the BDQ-saturated specimen had the highest overall resolutions and were used for building atomic models. To model the F_1_ region, the crystal structure of the *M. smegmatis* F_1_-ATPase containing subunits α_3_β_3_γε (PDB 6FOC) (Zhang et al., 2019) was fit rigidly into the maps with *UCSF Chimera* (Pettersen et al., 2004). Manual adjustment and building of the missing regions was performed in *Coot* (Emsley and Cowtan, 2004). To model the F_O_ region, the crystal structure of the *M. phlei* c ring in complex with BDQ (PDB 4V1F) (Preiss et al., 2015) was rigidly fit into the maps of the F_O_ regions from focused refinement. Manual adjustment of the model and building of subunit a and the membrane-embedded regions of subunits b-δ and b*′* were done in *Coot*. F_1_ and F_O_ models were then modified in *ISOLDE* (Croll, 2018) to improve dihedral angles and rotamer fitting before being refined in *Phenix* (Adams et al., 2010) with Ramachandran, rotamer and secondary structure restraints. One of the three N-terminal α helices from subunit α in the BDQ-free model was modeled as polyalanine (Table S4). Model quality was evaluated with *Molprobity* (Chen et al., 2010) and *EMRinger* (Barad et al., 2015). Full models of the BDQ-saturated rotational state 3 and BDQ-free rotational state 1 were prepared by rigid fitting of the F_1_ and F_O_ models into the maps with *UCSF Chimera*, manual adjustment of the connecting regions with *Coot*, and optimization of geometry with *ISOLDE*. Backbone models of the BDQ-saturated States 1 and 2 were built by flexible fitting of the F_1_ region of the BDQ-saturated rotational state 3 model into the corresponding maps with *ISOLDE*. The F_1_ regions were then combined with the ab*′*b-δ subcomplex and c ring from the BDQ-saturated rotational state 3 model fit separately into the maps as rigid bodies. Backbone models of the BDQ-free States 2 and 3 were built from the BDQ-free State 1 model in the same way. Figures and videos were generated with UCSF Chimera (Pettersen et al., 2004) and ChimeraX (Goddard et al., 2018).

---

**Fig. S1.**
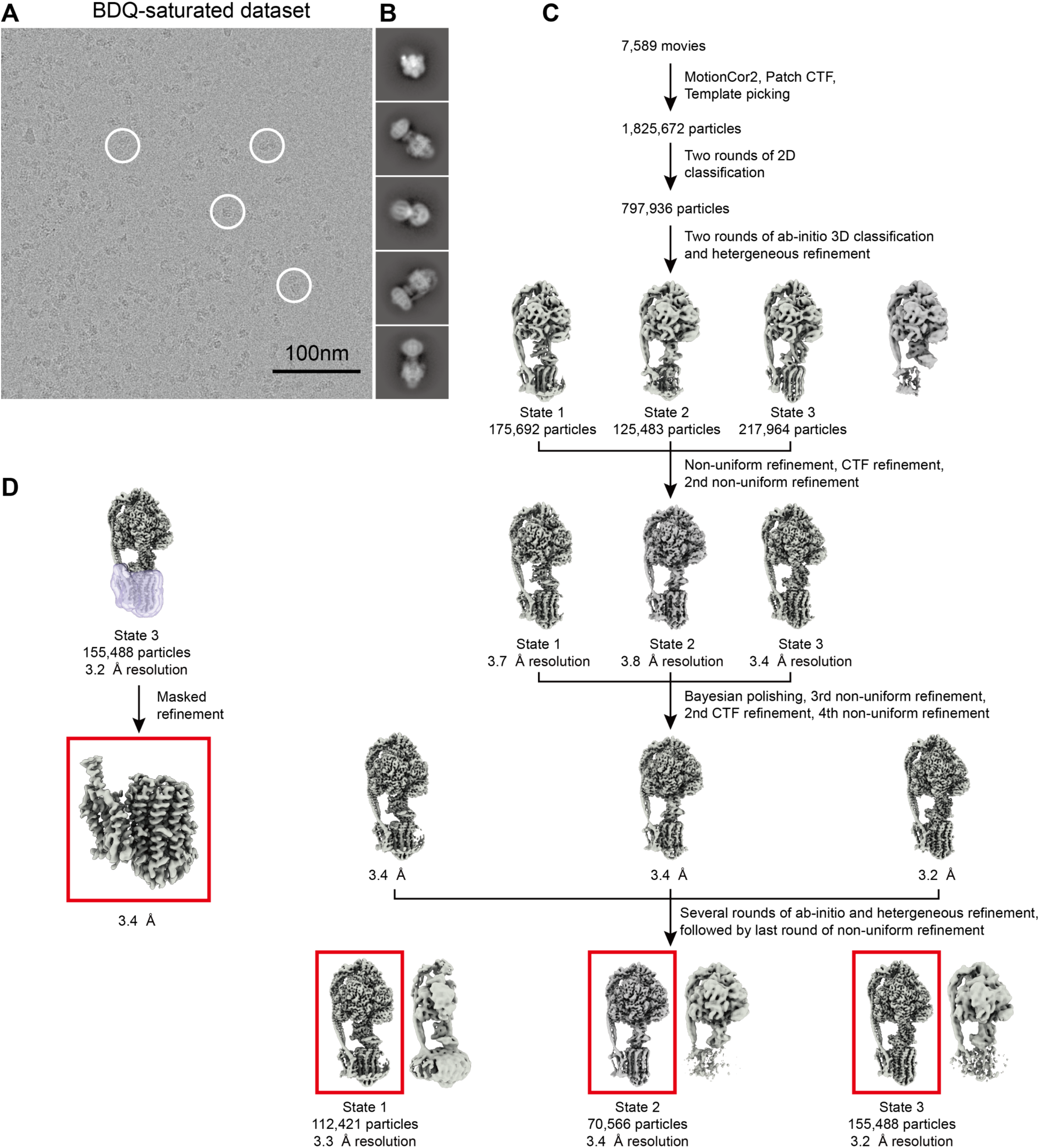
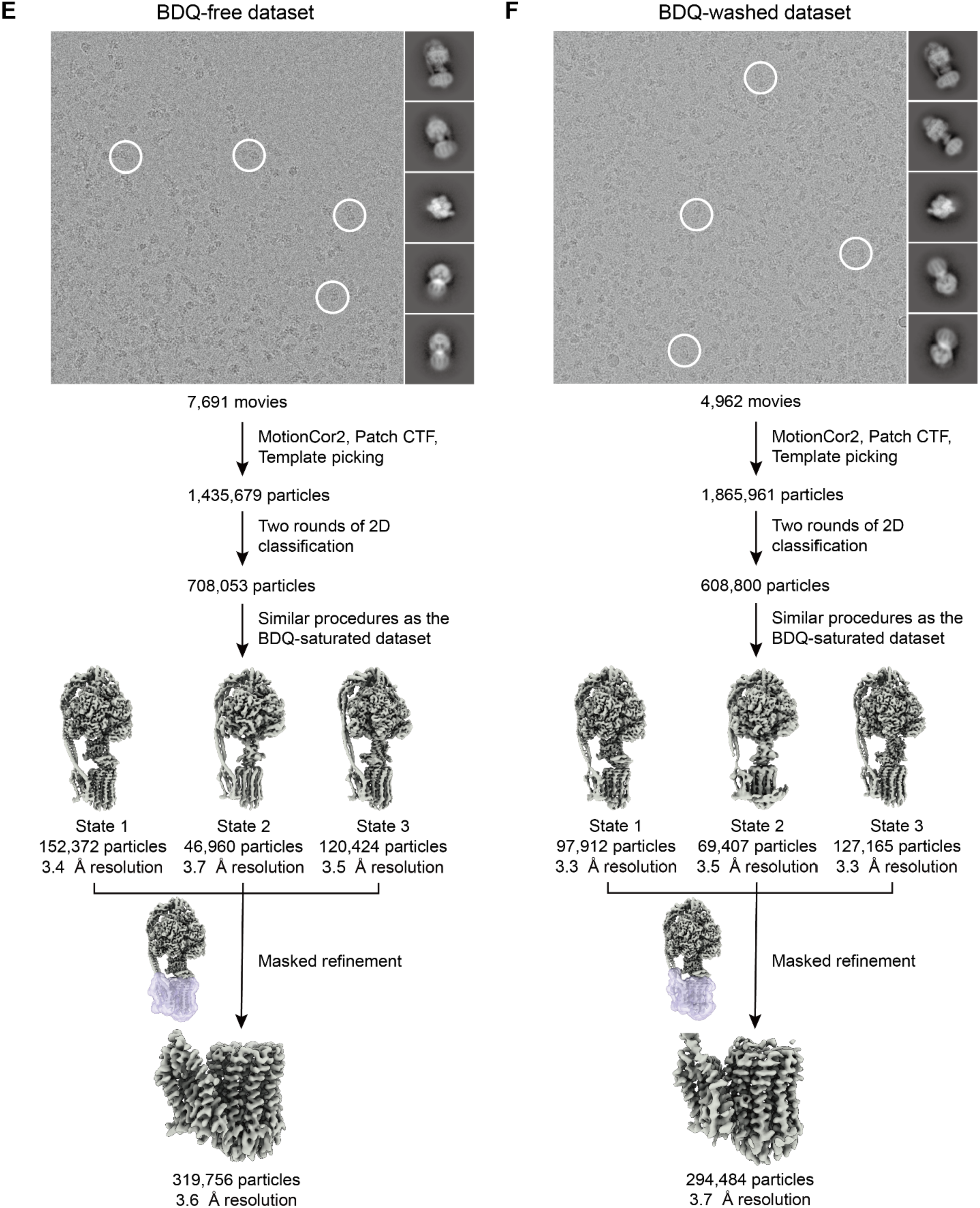
Workflow for cryoEM image analysis. Example micrograph (**A**) and 2D class average images (**B**) for BDQ-saturated *M. smegmatis* ATP synthase. Example particles are circled in white. Workflow for obtaining maps of the three rotational states (**C**) and F_O_ region (**D**). Summary of workflows for the BDQ-free (**E**) and BDQ-washed (**F**) datasets.

**Fig. S2.**
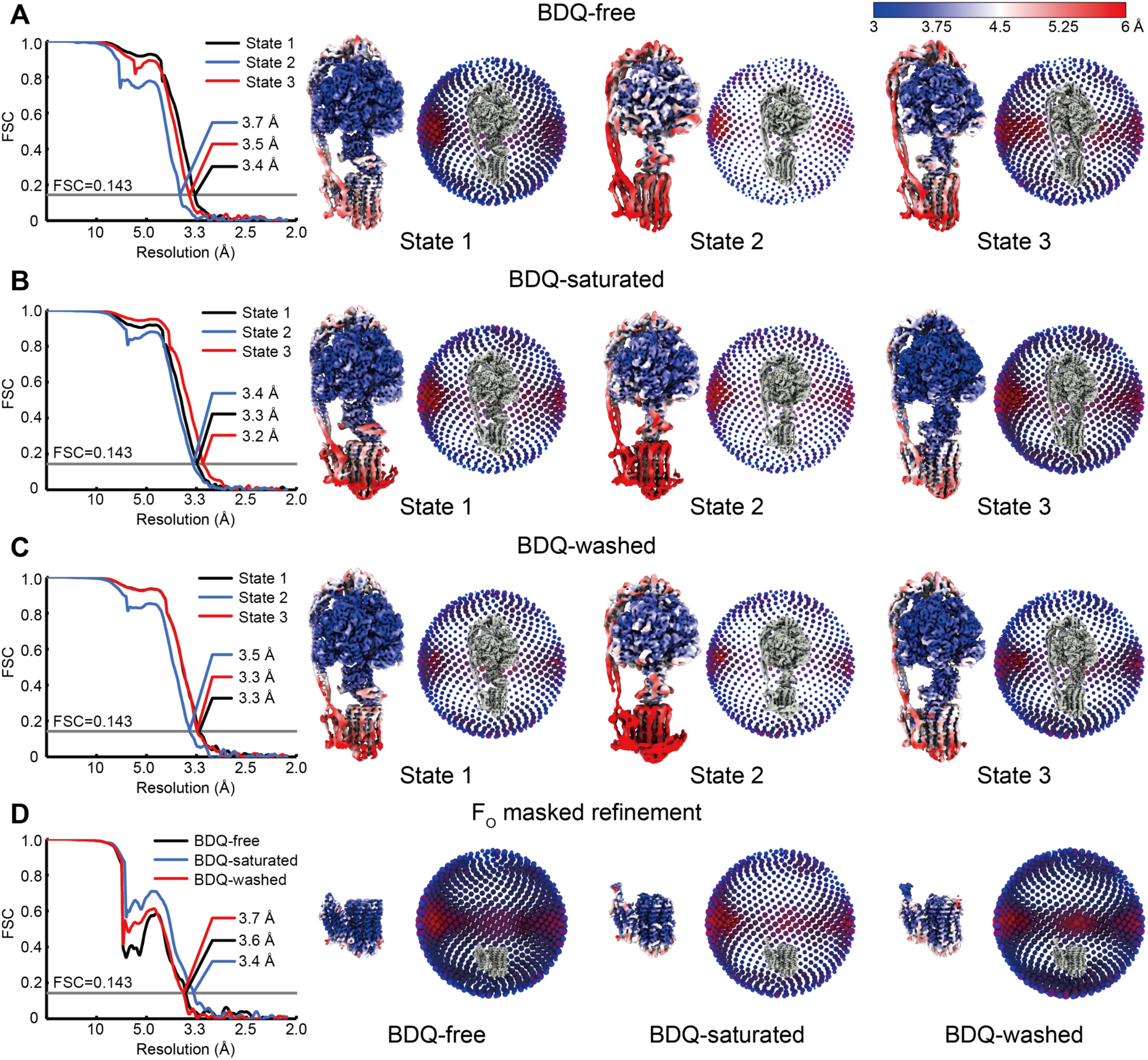
CryoEM map validation. Corrected Fourier shell correlation curves after gold-standard refinement, orientation distribution plots, and local resolution maps are shown for maps of the entire complex from the BDQ-free dataset (**A**), BDQ-saturated dataset (**B**), BDQ-washed dataset (**C**), as well as F_O_ region maps from focused refinement (**D**).

**Fig. S3.**
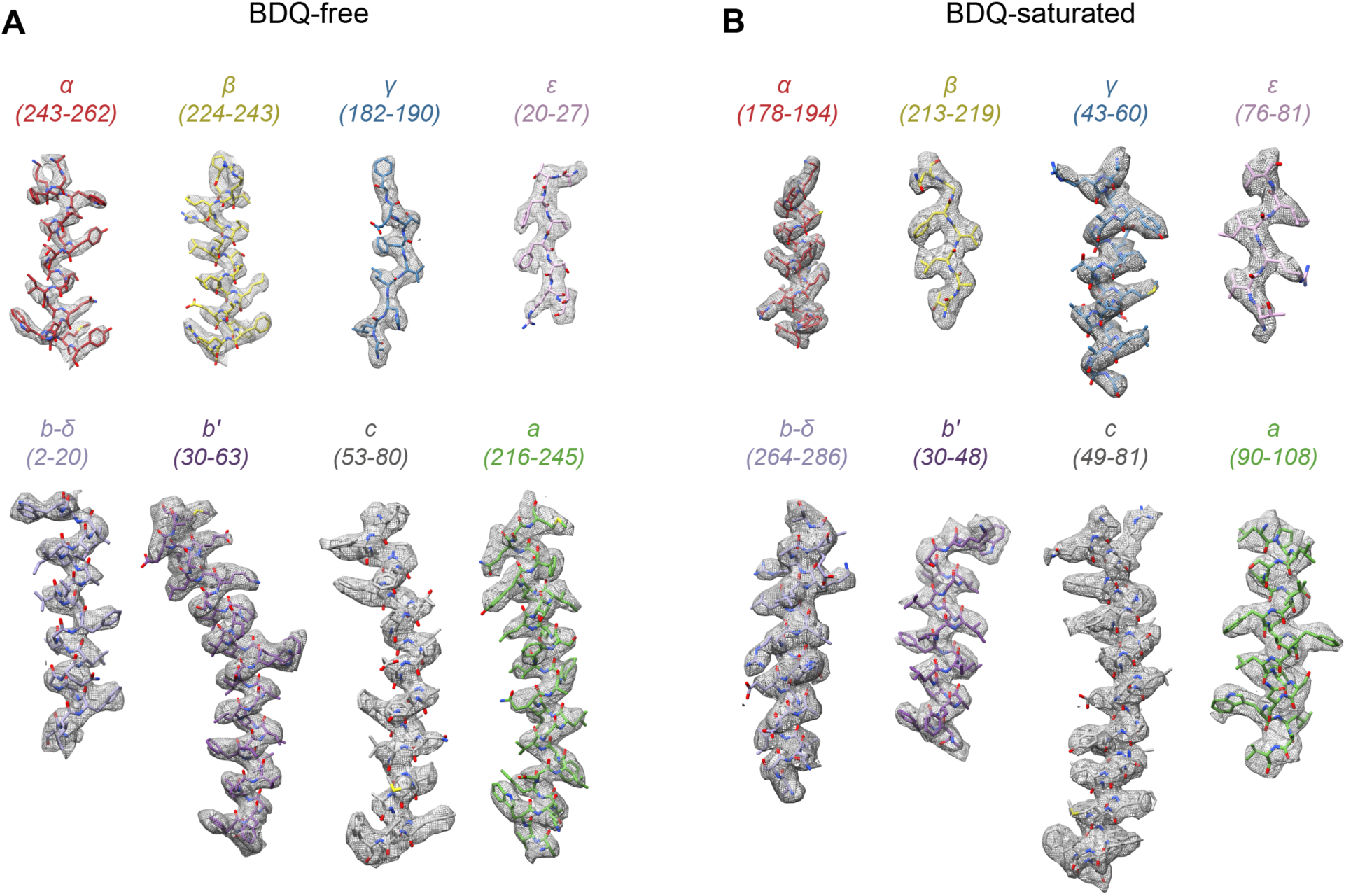
Examples of model-in-map fit to demonstrate map quality. Examples are shown from each subunit in the BDQ-free model (**A**) and BDQ-saturated model (**B**).

**Fig. S4.**
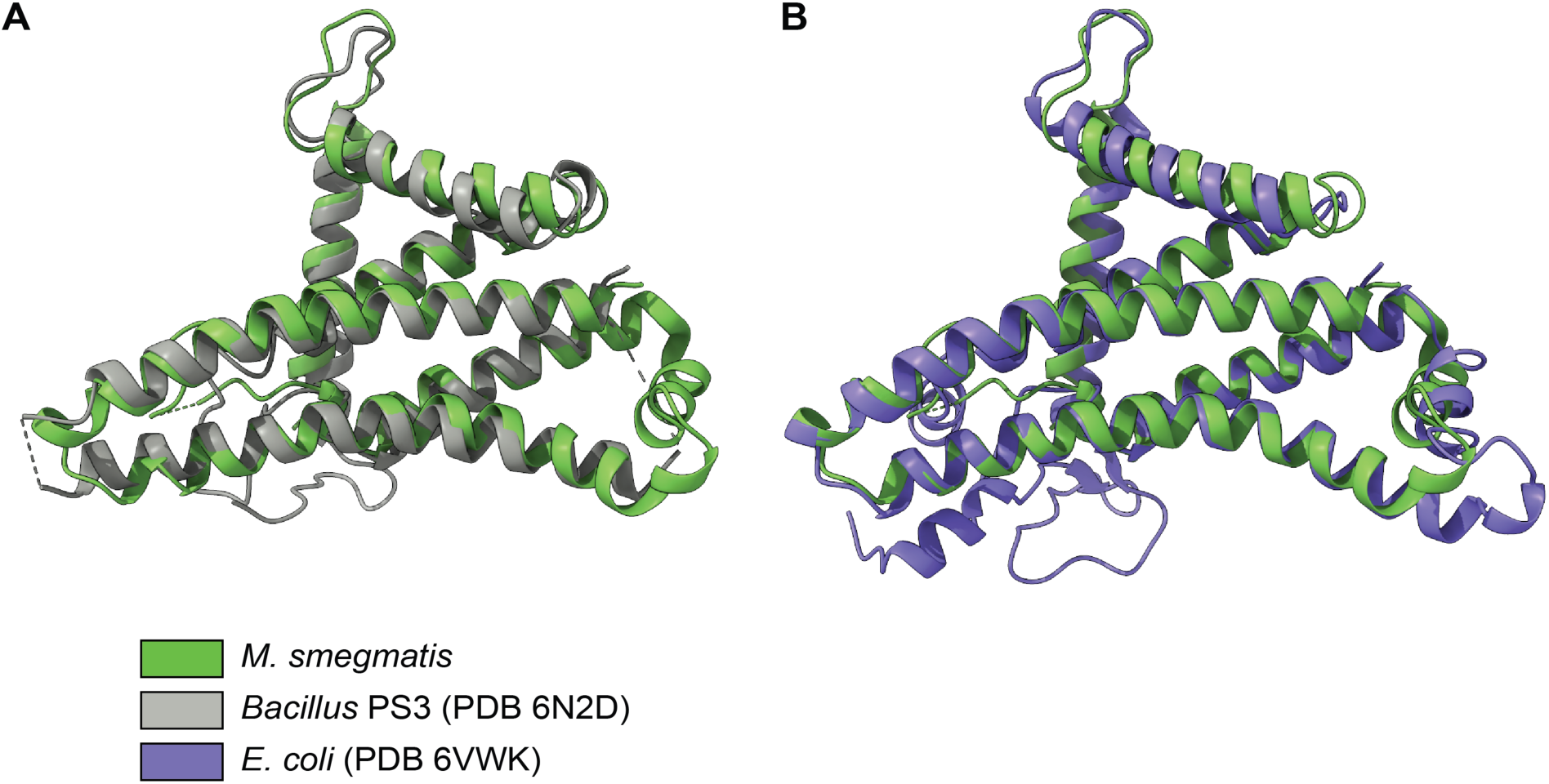
Comparison of the subunit a model from *M. smegmatis, Bacillus* PS3, and *E. coli*. **A**, The *Bacillus* PS3 structure is in grey and the *M. smegmatis* structure is in green. **B**, The *E. coli* structure is in purple and the *M. smegmatis* structure is in green.

**Fig. S5.**
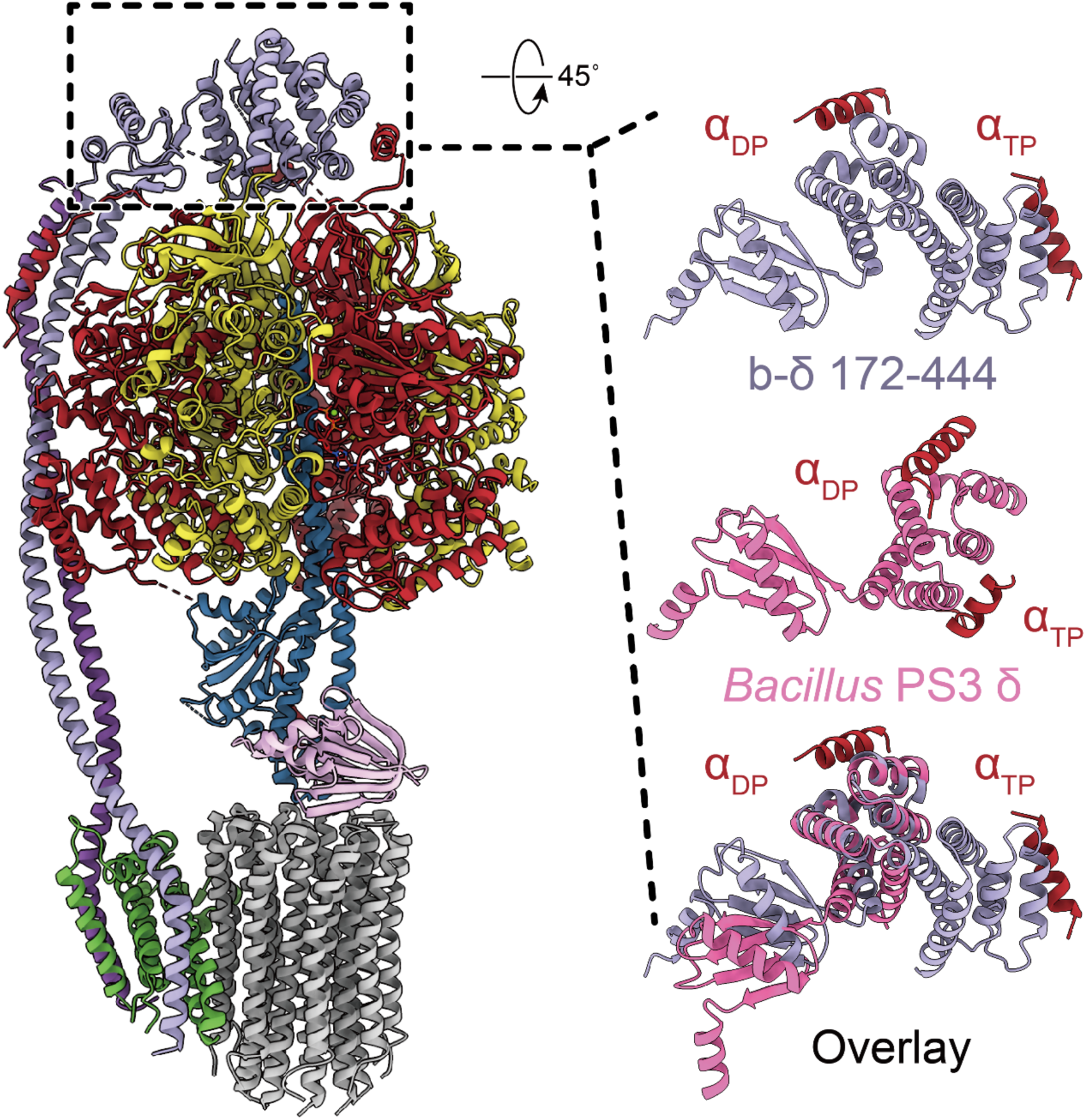
The N-terminal domain of the δ region of the b-δ fusion protein (*right side, top*) is substantially larger than the N-terminal domain of *Bacillus* PS3 δ subunit (*right side, middle*), composed of ∼270 residues, rather than ∼170 residues. The extra sequence contributes five additional α helices to the six found in the *Bacillus* PS3 δ subunit for a total of eleven α helices in the N-terminal domain of b-δ (Fig. 1B, *bottom*). *M. smegmatis* b-δ and *Bacillus* PS3 δ also form different interactions with the N-terminal sequences of the α subunits that help attach the peripheral stalk to F_1_ region (*right side, red*).

**Fig. S6.**
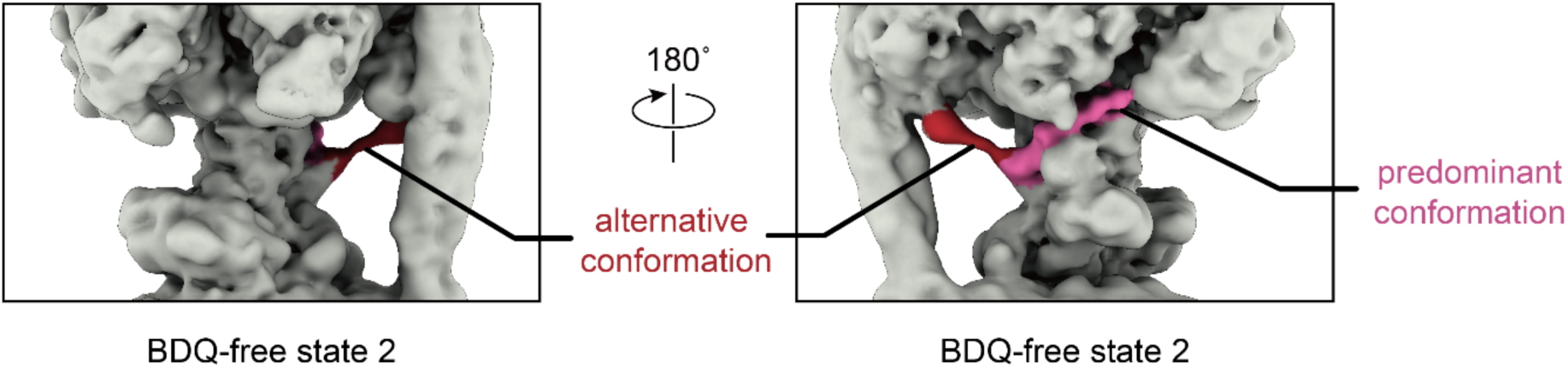
Two conformations of the α-extension are seen in rotational state 2. Density for the α-extension in its predominant conformation is colored pink while the alternative conformation is colored red.

**Fig. S7.**
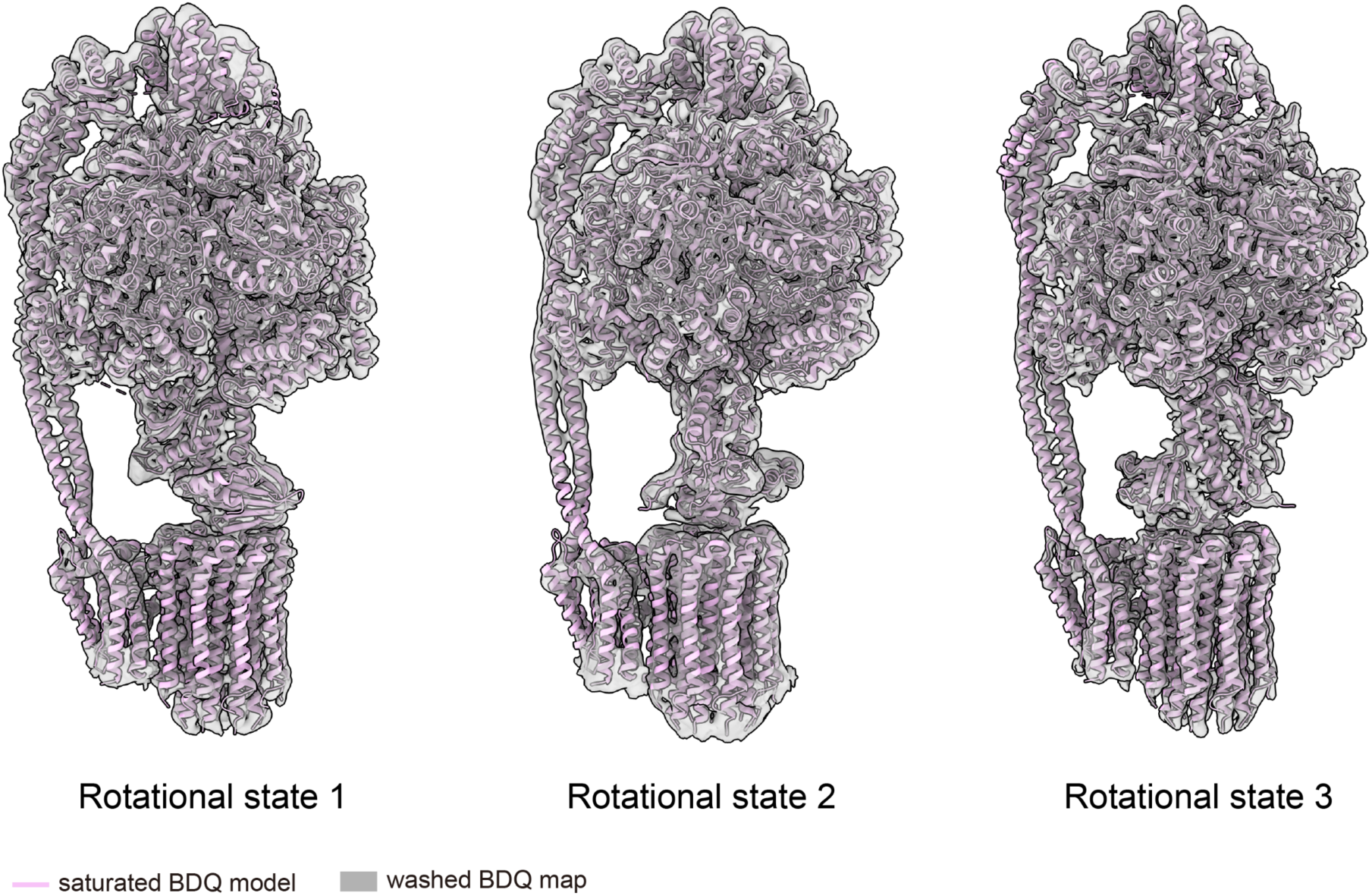
The BDQ-washed structures are in the BDQ-saturated conformation. Fit of the BDQ-saturated protein models in the BDQ-washed maps shows that the two preparations adopt the same conformation.

**Table S1.** EM statistics.

|  | BDQ-free | BDQ-saturated | BDQ-washed |
| --- | --- | --- | --- |
| **Data Collection** |  |  |  |
| Electron Microscope | Titan Krios G3 |  |  |
| Camera | Falcon 4 |  |  |
| Mode | Counting |  |  |
| Voltage (kV) | 300 |  |  |
| Nominal Magnification | 75,000× |  |  |
| Calibrated physical pixel size (Å) | 1.03 |  |  |
| Exposure time (s) | 9.6 | 8.9 | 8.9 |
| Total exposure (e<sup>-</sup>/Å<sup>2</sup>) | 45 | 41 | 41 |
| Exposure rate (e<sup>-</sup>/pixel/s) | 5 | 4.9 | 4.9 |
| Number of frames | 30 | 29 | 29 |
| Defocus range (μm) | 0.8-2.3 | 0.6-3.0 | 0.5-4.0 |
| **Image Processing** |  |  |  |
| Micrograph motion correction software | MotionCor2 |  |  |
| CTF estimation software | cryoSPARC v2 |  |  |
| Particle selection software | cryoSPARC v2 |  |  |
| 3D classification and refinement software | cryoSPARC v2 |  |  |
| Particle motion correction software | Relion 3 |  |  |
| Number of micrographs | 7,691 | 7,589 | 4,962 |
| Number of particle images selected | 1,435,679 | 1,825,672 | 1,865,961 |
| Number of particle images after cleanup | 708,053 | 797,936 | 608,800 |
| Number of particle images in final maps | 319,756 | 338,475 | 294,484 |
| Symmetry | C1 |  |  |
| **Model building** |  |  |  |
| Modeling software | Coot, ISOLDE |  |  |
| Refinement software | Phenix |  |  |

**Table S2.** Map statistics.

| Map | Resolution (Å,<br>FSC =0.143) | Applied B-factors (Å<sup>2</sup>) | Number of particle<br>images |
| --- | --- | --- | --- |
| BDQ-free state 1 | 3.4 | -114.6 | 152,372 |
| BDQ-free state 2 | 3.7 | -109.8 | 46,960 |
| BDQ-free state 3 | 3.5 | -94.7 | 120,424 |
| BDQ-saturated state 1 | 3.3 | -103.9 | 112,421 |
| BDQ-saturated state 2 | 3.4 | -100.5 | 70,566 |
| BDQ-saturated state 3 | 3.2 | -110.8 | 155,488 |
| BDQ-washed state 1 | 3.3 | -109.5 | 97,912 |
| BDQ-washed state 2 | 3.5 | -112.3 | 69,407 |
| BDQ-washed state 3 | 3.3 | -104.9 | 127,165 |
| BDQ-free focused Fo | 3.5 | -96.6 | 319,758 |
| BDQ-saturated focused Fo | 3.4 | -79.0 | 155,488 |
| BDQ-washed focused Fo | 3.7 | -90.7 | 294,484 |

**Table S3.** Atomic model statistics.

|  | **BDQ-free**<br>**state 1** | **BDQ-saturated**<br>**state 3** | **BDQ-free masked**<br>**Fo** | **BDQ-saturated**<br>**masked Fo** |
| --- | --- | --- | --- | --- |
| Number of residues | 4,830 | 4830 | 1022 | 1017 |
| Ligand | 3 Mg-ATP<br>1 Mg-ADP<br>1 Pi | 3 Mg-ATP<br>1 Mg-ADP<br>1 Pi<br>7 BDQ | - | 7 BDQ |
| Map-model CC_mask | 0.69 | 0.71 | 0.65 | 0.68 |
| RMS bond length (Å) | 0.007 | 0.007 | 0.007 | 0.008 |
| RMS bond angle (°) | 1.027 | 1.055 | 0.642 | 0.736 |
| Clash score | 3.15 | 3.75 | 3.73 | 5.86 |
| **Ramachandran** |  |  |  |  |
| Favoured (%) | 98.36 | 97.80 | 98.09 | 97.58 |
| Allowed (%) | 1.51 | 2.05 | 1.91 | 2.42 |
| Outliers (%) | 0.13 | 0.15 | 0.00 | 0.00 |
| Rotamer outliers (%) | 0.14 | 0.14 | 0.45 | 0.15 |
| C<sup>β</sup> outliers | 0.00 | 0.00 | 0.00 | 0.00 |
| CaBLAM outliers (%) | 0.98 | 0.87 | 0.41 | 0.62 |
| **Average B factors (Å<sup>2</sup>)** |  |  |  |  |
| Protein | 67.94 | 83.37 | 22.48 | 63.71 |
| Ligands | 68.92 | 69.99 | - | 65.71 |
| MolProbity score | 1.11 | 1.21 | 1.16 | 1.41 |
| EMRinger score | 2.60 | 3.16 | 2.65 | 3.40 |

**Table S4.** Residues included in atomic models. F_O_ atomic models contain the same residues as the F_O_ region in the corresponding models of the full structure.

| Chain ID (subunit) | BDQ-free state 1 |  | BDQ-saturated state 3 |
| --- | --- | --- | --- |
|  | Atomic model | Poly alanine | Atomic model |
| A ($\alpha_1$) | 5-516, 533-545 | - | 7-520 |
| B ($\alpha_2$) | 11-22, 28-520 | - | 8-22, 28-520 |
| C ($\alpha_3$) | 29-520 | 11 residues at the N terminus | 9-22, 29-515, 533-546 |
| D ($\beta_1$) | 8-471 | - | 8-471 |
| E ($\beta_2$) | 8-471 | - | 8-471 |
| F ($\beta_3$) | 8-471 | - | 8-471 |
| G ($\gamma$) | 4-164, 178-213, 222-303 | - | 4-164, 178-213, 222-303 |
| H ($\epsilon$) | 3-119 | - | 3-119 |
| a (a) | 31-113, 123-246 | - | 31-113, 123-246 |
| b (b') | 20-164 | - | 24-164 |
| d (b-δ) | 1-165, 172-255, 260-285, 288-331, 337-444 | - | 1-157, 169-331, 337-444 |
| 1-9 (c<sub>1</sub>-c<sub>9</sub>) | 5-85 | - | 5-85 |

**Table S5.** Deposited maps and models.

| Map | EMDB code | PDB ID | Model type |
| --- | --- | --- | --- |
| BDQ-free state 1 | EMD-22311 | 7JG5 | atomic |
| BDQ-free state 2 | EMD-22312 | 7JG6 | backbone |
| BDQ-free state 3 | EMD-22313 | 7JG7 | backbone |
| BDQ-saturated state 1 | EMD-22314 | 7JG8 | backbone |
| BDQ-saturated state 2 | EMD-22315 | 7JG9 | atomic |
| BDQ-saturated state 3 | EMD-22316 | 7JGA | backbone |
| BDQ-washed state 1 | EMD-22317 | - | - |
| BDQ-washed state 2 | EMD-22318 | - | - |
| BDQ-washed state 3 | EMD-22319 | - | - |
| BDQ-free focused F<sub>O</sub> | EMD-22320 | 7JGB | atomic |
| BDQ-saturated focused F<sub>O</sub> | EMD-22321 | 7JGC | atomic |
| BDQ-washed focused F<sub>O</sub> | EMD-22322 | - | - |

**Video 1. Release of the inhibitory α extension during ATP synthesis**. The video shows interpolation between the different rotational states of the enzyme, including the alternative conformation of rotational state 2, which suggests the sequence of events that occur during release of the α-extension during ATP synthesis and binding of the α-extension to subunit γ to inhibit ATP hydrolysis. **Video may be viewed as a loop**.

**Video 2. Binding of BDQ to the mycobacterial ATP synthase**. The video shows interpolation between the conformation of the ATP synthase in the BDQ-free and BDQ-bound states. BDQ in the leading, c-only, and lagging binding sites are colored pink, yellow, and blue, respectively. Phe69 in subunit c and Phe221 in subunit a, which undergo large conformational changes upon drug binding, are shown as space-filling models. **Video may be viewed as a loop**.

